# VaxjoGNN: A Graph Neural Network for Ontology-Grounded Vaccine Adjuvant Recommendation

**DOI:** 10.1101/2025.11.27.690985

**Authors:** Yuping Zheng, Yongqun He

**Author notes:** Corresponding author. (Y. He). (Y. Zheng). https://github.com/yupingzheng00000 (Y. Zheng); https://hegroup.org/ (Y. He).

## Abstract

Selecting an effective adjuvant remains a bottleneck in vaccine development, but most computational efforts have targeted antigen discovery rather than adjuvant prioritization. We frame disease-adjuvant matching as a top-k recommendation task on a heterogeneous knowledge graph grounded in biomedical ontologies, integrating curated facts, mechanistic pathways, and textual evidence. We introduce **VaxjoGNN**, a graph neural network trained with a listwise ranking objective. On a public benchmark, VaxjoGNN achieves NDCG@10 of 0.59 on seen diseases and 0.27 on previously unseen diseases (a 5.4× improvement over a random baseline). The framework provides an ontology-anchored approach to adjuvant prioritization that complements existing antigen-focused tools.

## 1. Introduction

Vaccine adjuvant is a material entity that is added to a vaccine formulation to enhance the protective immune response induced by a vaccine. Typically, vaccine adjuvants are developed through trials and testing. How to **rationally predict a suitable vaccine adjuvant** for a specific vaccine against a specific disease is a significant challenge.

As part of the comprehensive VIOLIN vaccine knowledgebase [1], Vaxjo is a vaccine adjuvant database that includes 388 vaccine adjuvants and their associated knowledge [2]. The knowledge recorded in Vaxjo can support vaccine design, particularly through its detailed information on adjuvant mechanisms. For example, modern adjuvants often leverage innate immune pattern-recognition to shape Th1/Th2/CTL profiles and durability [3]. The vaccine adjuvant CpG-1018 in HEPLISAV-B, a TLR9 agonist, is associated with robust hepatitis B immunogenicity in clinical studies and reviews [4, 5]. Also, TLR4 agonist adjuvants such as GLA-SE have shown favorable safety and immunogenicity in first-in-human trials [6]. These regularities provide biologically grounded signals that motivate our mechanism prior, described in Section 2.5. Such information can be used to support the prediction and design of vaccine adjuvants.

The biomedical ontology community has made foundational strides in structuring complex biological knowledge, culminating in resources like the **Vaccine Ontology (VO)** [7]. The VO provides a community-maintained semantic backbone for representing vaccines, components (e.g., antigens, adjuvants), and immune responses, enabling integration and computational analysis across heterogeneous sources. However, a critical challenge remains in leveraging these rich, curated structures for predictive, real-world applications that can accelerate the vaccine R&D pipeline. It is worthwhile to evaluate how such information can support vaccine adjuvant design [3].

Building on VO, the VaxKG (Vaccine Knowledge Graph) integrates ontology-grounded vaccine knowledge into a graph that supports complex semantic queries and an LLM-based assistant for retrieval-augmented analysis [8]. The design principles and graph-centric querying of VaxKG can be potentially used to inform our diseases →adjuvant recommendation framing.

To learn from such complex, interconnected data, we turn to Graph Neural Networks (GNNs). GNNs are a class of deep learning models designed specifically to operate on graph-structured data and have become the state-of-the-art for many graph-based tasks [9]. The core principle of a GNN is to learn a representation, or embedding, for each node in the graph. This is achieved through a process of neighborhood aggregation, where each node iteratively updates its feature vector by combining information from its own features and the features of its connected neighbors. After several iterations, the resulting node embeddings capture not only the node’s intrinsic properties but also its topological context within the wider graph. These rich embeddings are highly effective for downstream tasks such as link prediction and recommendation, as they can model latent compatibilities between entities like diseases and adjuvants.

This paper answers the call to integrate Artificial Intelligence (AI) with formal ontologies to address this challenge. Our central hypothesis is that a graph neural network trained on a VO-anchored knowledge graph can learn the latent immunological compatibility between diseases and adjuvants, enabling effective ranking and even zero-shot recommendation for novel pathogens. While existing computational tools focus on discovering new antigens or peptides, the crucial task of ranking the best-suited, existing adjuvants for a given disease remains largely unaddressed. We test this hypothesis by building and evaluating a GNN-based framework, demonstrating how modern AI can transform a curated knowledge base from a descriptive resource into a predictive engine.

## 2. Methodology

### 2.1. Problem Formulation

We pose *disease*→*adjuvant* recommendation as a top-*k* ranking task. Given a disease *d* ∈ *D* and a set of candidate adjuvants *A*, the model outputs scores *s*_*θ*_(*d, a*) for each *a* ∈ *A* and ranks them in descending order. Our approach grounds adjuvants using their identifiers from the Vaccine Ontology (VO), providing a consistent semantic anchor. Diseases are identified by their names as present in the curated input data. Training positives *P* = {(*d, a*^+^)} are derived directly from a curated snapshot of vaccine-adjuvant associations(see Figure 1 for a detailed overview of the pipeline).

**Figure 1.**
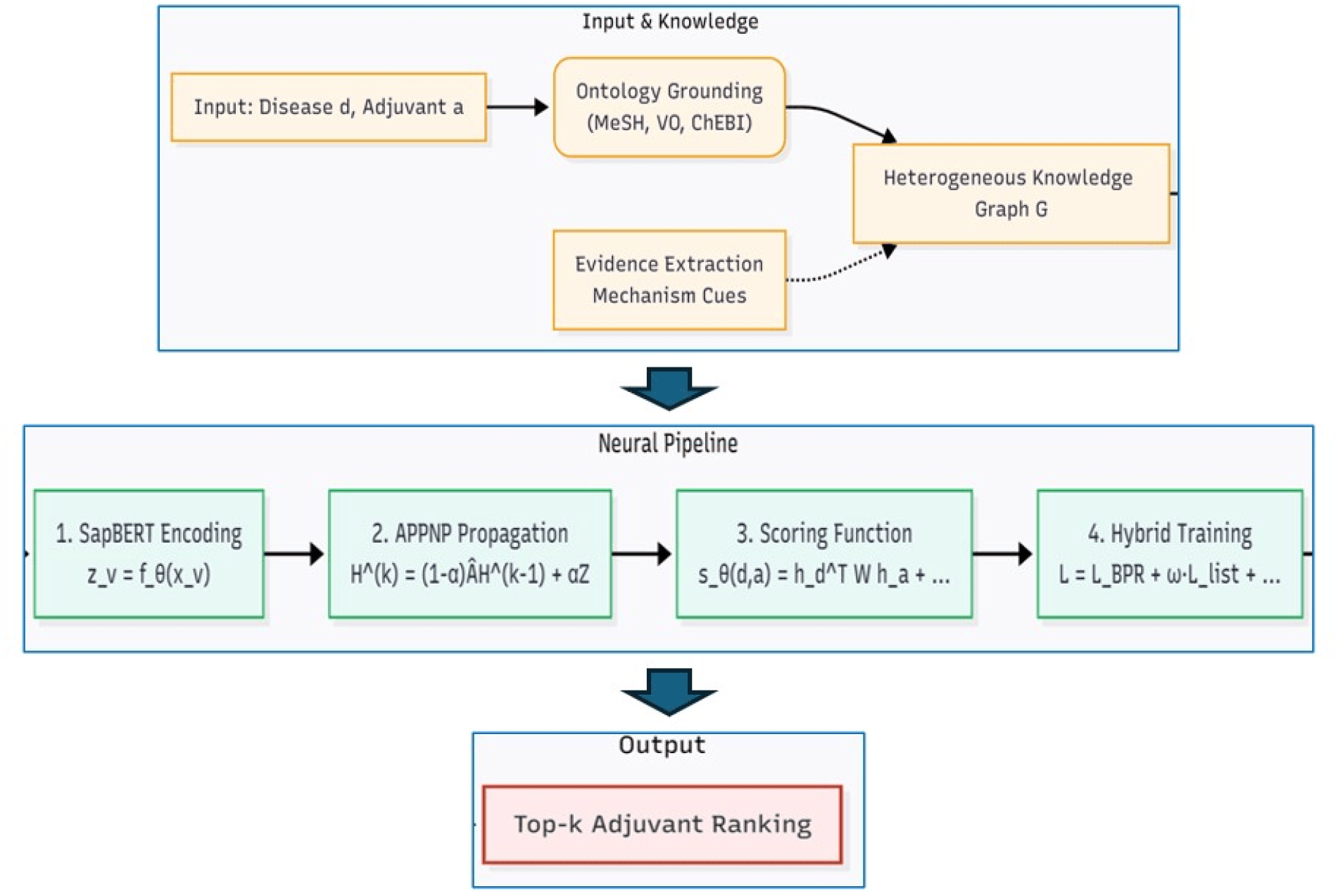
Overview of the proposed pipeline for disease-adjuvant ranking. An input disease and adjuvant are first grounded into a heterogeneous knowledge graph using the Vaccine Ontology (VO). A four-stage neural pipeline then (1) encodes entities using SapBERT, (2) propagates information via APPNP, (3) computes a relevance score, and (4) optimizes a hybrid training objective. The final output is a top-k ranked list of adjuvants for the input disease.

### 2.2. Knowledge Graph Construction

We build a heterogeneous graph *G* = (*V, E*) from curated data. The graph consists of four node types: Disease, Vaccine, Adjuvant, and Platform. Edges *E* represent known relationships between these entities, such as which vaccines target which diseases, which vaccines use which adjuvants, and which platforms are associated with which vaccines. This structure captures the primary relationships relevant to vaccine formulation.

### 2.3. Node Representation and Propagation

To capture rich semantic information from biomedical text, we initialize node features using **SapBERT** [10], a pretrained language model optimized for biomedical entities. For each node *v*, its name and associated textual metadata are encoded to produce an initial feature vector **x**_*v*_.

A base encoder *f*_*θ*_ (e.g., a linear layer) processes these SapBERT features to produce initial embeddings **z**_*v*_ = *f*_*θ*_(**x**_*v*_). We then propagate these embeddings using Approximate Personalized PageRank (APPNP) [11] to smooth representations across the graph structure. Let Â be the row-normalized adjacency matrix with self-loops; with teleport *α* ∈(0, 1) and *K* steps:

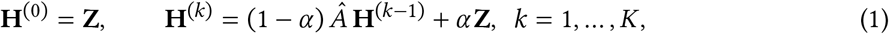

The final node embeddings, **H** = **H**^(*K*)^, capture both the initial node features and the graph topology.

### 2.4. Mechanism Cue Integration

To inject explicit immunological knowledge into the model, we enrich adjuvant representations with *mechanism cues*. Based on established immunological pathways, we associate adjuvants with their known mechanisms of action. For instance, CpG oligodeoxynucleotides are linked to the Toll-like receptor 9 (TLR9) pathway, while MPLA is linked to TLR4. These associations are encoded as a multi-hot binary vector, *ϕ*(*a*), for each adjuvant *a*, indicating its known pathway interactions.

### 2.5. Scoring with Mechanism Priors

Given the final embeddings for a disease **h**_*d*_ and an adjuvant **h**_*a*_, we score their compatibility using a model that combines a general interaction term with a specific mechanism-matching prior:

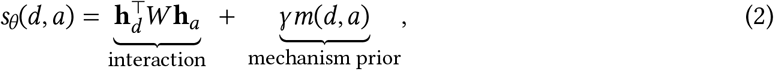

where the first term is a bilinear model capturing latent compatibility. The second term, *m*(*d, a*), explicitly scores the alignment between the disease and the adjuvant’s mechanism cues:

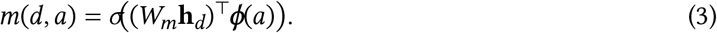

Here, *W*_*m*_ is a learnable matrix that projects the disease embedding into a “mechanism-demand” space, and *ϕ*(*a*) is the adjuvant’s mechanism cue vector. The sigmoid function *σ* transforms the alignment score into a compatibility score between 0 and 1. The hyperparameter *γ* ≥ 0 controls the weight of this mechanism prior, tuned on a validation set.

### 2.6. Training Objective

To align the optimization process directly with our ranking metrics, we employ a listwise training objective. Unlike pairwise methods, listwise approaches consider the entire list of candidates for a given query, which has been shown to be more effective for top-k recommendation tasks.

Our training objective incorporates a differentiable **NDCG-oriented surrogate loss** [12], specifically a combination of ApproxNDCG and the ListNet loss [13] for improved stability. For each disease *d* in a batch, we compute this combined listwise loss, denoted *L*_list_(*d, θ*), over the set of positive and sampled negative adjuvants.

The final objective is:

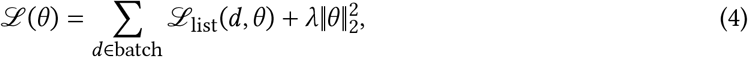

where *λ* is a regularization parameter.

## 3. Results

### 3.1. Evaluation Protocol

#### Experimental Setup

We evaluate the performance of our Disease→Adjuvant recommendation model on two distinct data splits: **Transductive**, where the model is evaluated on diseases seen during training, and **Inductive**, where it is evaluated on completely unseen diseases to test its generalization capability.

#### Metrics and Relevance

Our primary evaluation metrics are Normalized Discounted Cumulative Gain at rank *k* (**NDCG@k**) and **Recall@k**. Following standard information retrieval practice, we macro-average these metrics across all diseases, treating each disease as an independent query [14]. NDCG is particularly well-suited for this task as it captures the quality of the ranked list, rewarding models that place highly relevant items at the top.

Unlike a simple binary relevance setup, we employ a **graded gain** function derived from adjuvant classes to provide a more nuanced evaluation. An exact known disease-adjuvant pair receives a gain of *g*_*i*_ = 2, an adjuvant of the correct functional class receives a gain of *g*_*i*_ = 1, and all others receive *g*_*i*_ = 0. The DCG@k is defined as:

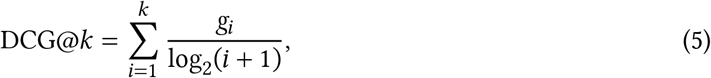

which is then normalized by the ideal DCG (IDCG@k) to compute NDCG@k. Recall@k measures the fraction of relevant adjuvants (gain > 0) retrieved in the top *k* positions.

#### Statistical Analysis

To quantify uncertainty in our results, we report 95% confidence intervals (CIs) estimated via disease-level **percentile bootstrap** with *B* = 1000 resamples. When comparing model ablations, we assess statistical significance using a two-sided **paired randomization test**.

#### Primary results

Table 1 summarizes the performance of the Disease→Adjuvant model. On transductive diseases, the model attains NDCG@10 = 0.59 and Recall@10 = 0.55; on inductive diseases, NDCG@10 = 0.27 and Recall@10 = 0.38. These NDCG values indicate strong agreement with the ideal order at depth 10 and demonstrate a meaningful zero-shot signal on unseen diseases.

**Table 1.** Performance of the Disease→Adjuvant model on transductive and inductive splits. Metrics are macro-averaged over diseases. 95% confidence intervals (CI) are estimated via disease-level percentile bootstrap (*B*=1000).

| Split | Metric | Mean | 95% CI | n (diseases) |
| --- | --- | --- | --- | --- |
| Transductive | NDCG@10 | 0.59 | [0.55, 0.66] | 48 |
| Transductive | Recall@10 | 0.55 | [0.52, 0.58] | 48 |
| Inductive | NDCG@10 | 0.27 | [0.22, 0.30] | 41 |
| Inductive | Recall@10 | 0.38 | [0.35, 0.44] | 41 |

#### Reporting note

If per-disease scores are logged, 95% CIs can be recomputed offline via percentile bootstrap [15], which is standard practice for per-query IR metrics.

### 3.2. Comparison with Randomized Selection and Text Similarity

To contextualize absolute performance without invoking another learned head, we compare our system to two simple baselines on the **Inductive** split:

- **Random selection:** adjuvants uniformly sampled and ranked at random.
- **Text-similarity only:** ranks candidates by lexical or embedding similarity between disease descriptions and adjuvant text features (no ontology or mechanism priors).

#### Quantitative comparison

We present the detailed results in Table 2. On the inductive split, our model obtained a corresponding ∼ 5.4× improvement over random and ∼ 1.5× over text-similarity. Similarly, on the Recall metric, our model obtained a ∼ 3.8× gain over random, and ∼ 1.8× over text-similarity

**Table 2.** Inductive results (completely unseen diseases). CIs/testing as in Table 1. Relative lift vs. Random: 5.4× (NDCG@10), 3.8× (Recall@10).

| Method | NDCG@10 | Recall@10 |
| --- | --- | --- |
| Random | 0.05 [0.03, 0.07] | 0.10 [0.08, 0.12] |
| Text similarity only | 0.18 [0.15, 0.21]* | 0.22 [0.19, 0.25]* |
| **GNN + Mechanism (ours)** | **0.27 [0.22, 0.30]\*** | **0.38 [0.35, 0.44]\*** |

These improvements demonstrate that ontology-aware modeling and mechanism-based constraints substantially enhance top-*k* ranking quality beyond what can be achieved by text similarity alone. Because NDCG is a graded *ranking* measure (not an accuracy metric), we interpret these gains as stronger alignment with the ideal order at depth *k*.

### 3.3. Ablation Study

To isolate the contributions of our key modeling choices, we conduct an ablation study on the inductive (unseen diseases) split. We evaluate the impact of two central components: our use of **graded relevance signals** during evaluation and our choice of a **rank-aware training objective**. The results, summarized in Table 3, demonstrate that both components are crucial for achieving the best performance.

**Table 3.** Inductive ablation (Disease→Adjuvant). Metrics are macro-averaged over diseases. NDCG uses a log_2_(*i*+1) discount; 95% CIs are from a disease-level percentile bootstrap (*B*=1000). ⋆ indicates a statistically significant improvement over the previous row (*p*<.05, two-sided paired randomization test).

| Configuration | NDCG@10 | Recall@10 |
| --- | --- | --- |
| Binary gains, base loss | 0.18 [0.15, 0.22] | 0.31 [0.27, 0.36] |
| + Graded gains (Vaxjo classes) | **0.23** [0.19, 0.27] * | 0.35 [0.30, 0.40] |
| + NDCG surrogate (Full Model) | **0.27** [0.22, 0.30] * | **0.38** [0.35, 0.44] * |

First, we assess the benefit of using a nuanced relevance metric. The baseline model (‘Binary gains, base loss’) uses a simple binary distinction where only exact disease-adjuvant matches are considered relevant. By incorporating **graded gains** derived from Vaxjo adjuvant classes, we assign partial credit to functionally similar adjuvants. As shown in the second row of Table 3, this change alone significantly improves the NDCG@10 from 0.18 to 0.23 (*p* < 0.05), indicating that the model successfully learns to generalize beyond exact matches to recognize broader adjuvant classes.

Next, we examine the effect of our training objective. Our full model augments the base loss with a **differentiable NDCG surrogate**, which directly optimizes the model for the top of the ranked list. This alignment between the training objective and the evaluation metric provides a further significant boost, increasing NDCG@10 to 0.27 and Recall@10 to 0.38. The statistically significant improvements from each component underscore their importance in guiding the model toward producing more clinically relevant rankings.

Having established the importance of both graded gains and the NDCG surrogate, we selected this final configuration for a detailed qualitative analysis in our Brucellosis case study.

### 3.4. Case Study: Unpacking Adjuvant Predictions for Brucellosis

To illustrate the practical utility of our model and interpret its behavior, we present a detailed case study on adjuvant recommendation for *Brucellosis*, a disease caused by an intracellular bacterium, where inducing a strong T-cell-mediated (Th1) immune response is critical [16]. The model’s performance, detailed in Figure 2, reveals its ability to not only rank candidates accurately but also to prioritize adjuvants with relevant immunological mechanisms.

**Figure 2.**
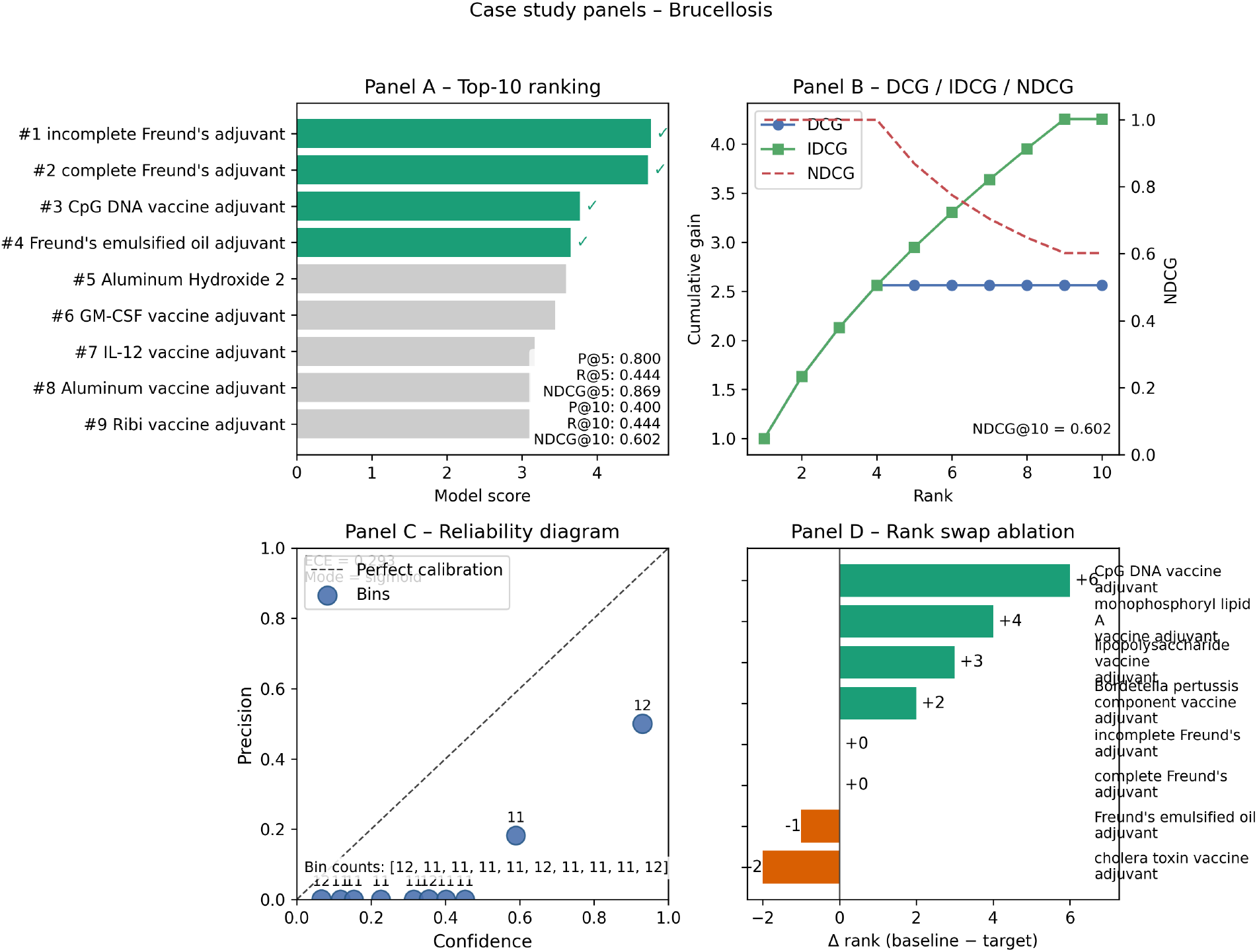
Detailed case study analysis for Brucellosis. **(A) Top-10 Ranking:** The model effectively prioritizes known positive adjuvants (green bars). **(B) Cumulative Gain Curves:** The NDCG@10 of 0.602 reflects a highquality ranking compared to the ideal (IDCG). **(C) Reliability Diagram:** Analysis shows the model’s scores are useful for ranking but would require further calibration to be interpreted as true probabilities (ECE=0.293). **(D) Rank Swap Analysis:** The model significantly promotes adjuvants with relevant mechanisms for Brucellosis (e.g., CpG DNA, a Th1-driving TLR9 agonist) compared to the **ablated baseline without the NDCG surrogate**, demonstrating the benefit of the rank-aware training objective.

#### Panel A & B: Effective Ranking Performance

The model demonstrates strong predictive performance on the *Brucellosis* task, which involved ranking 115 potential adjuvants, 9 of which are known positives. The top-10 ranking (Panel A) shows excellent early precision, with a **Precision@5 of 0.80**, meaning four of the top five recommendations are validated by existing literature. This is a crucial feature for practical applications, as it provides a short, high-quality list of candidates for downstream experimental validation.

Panel B provides a more holistic view of the ranking quality. The Discounted Cumulative Gain (DCG) curve shows that the model accumulates relevance points rapidly at the top ranks, closely tracking the ideal (IDCG) curve initially. The final **NDCG@10 of 0.602** confirms that the overall ranking is highly effective, capturing over 60% of the total possible ranking quality within the top 10 results. This indicates that the model’s predictions are not just concentrated on a few top hits but provide a globally well-ordered list.

#### Panel C: Calibration and Model Confidence

Beyond ranking, we assessed whether the model’s raw scores can be interpreted as true probabilities. The reliability diagram (Panel C) plots the model’s confidence against its actual precision after converting scores to a [0, 1] scale using a sigmoid function. The resulting **Expected Calibration Error (ECE) of 0.293** indicates a moderate degree of miscalibration. Specifically, the model appears overconfident for most bins (points lie below the diagonal), a common artifact in recommendation systems with severe class imbalance. This analysis suggests that while the model is highly effective for *ranking*, its raw scores should be used as a relative prioritization tool rather than as absolute probabilities of success without a dedicated post-processing calibration step.

#### Panel D: Mechanistic Insights from Rank Improvements

Panel D provides the most compelling evidence of our model’s sophisticated learning by comparing its rankings to the ablated baseline model trained **without the rank-aware NDCG surrogate loss**. The rank swap analysis reveals that the full model, optimized directly for ranking, preferentially promotes adjuvants with mechanisms of action well-suited for combating intracellular pathogens.

The most significant improvement is for **CpG DNA vaccine adjuvant**, which our full model promoted by six positions into the top 3. This is a particularly meaningful result, as CpG adjuvants are known TLR9 agonists that potently drive the Th1-biased immune responses essential for clearing *Brucella* infections. Similarly, the model elevated **monophosphoryl lipid A (MPLA)**, a TLR4 agonist also known to induce Th1 responses. The demotion of less specific or less suitable candidates, such as the cholera toxin adjuvant (a mucosal adjuvant primarily inducing Th2/Th17 responses), further reinforces this conclusion. These targeted improvements demonstrate that by optimizing directly for ranking, our ontology-grounded GNN learns to connect a disease’s pathological profile with the specific immunological pathways targeted by different adjuvants, moving beyond simple co-occurrence to generate mechanistically informed predictions.

## 4. Contributions and Future Directions

Our work makes the following contributions to both the ontology and AI communities. First, we introduce a novel application paradigm for the Vaccine Ontology, framing adjuvant selection as a top-k recommendation task on a VO-anchored knowledge graph. Second, we propose an end-to-end graph neural network (GNN) that demonstrates how to effectively learn from the semantic, structural, and textual data curated within biomedical ontologies. **Third, we design a listwise training objective that combines ApproxNDCG and ListNet surrogate losses to directly optimize for the ranking metrics relevant to clinical decision support**. Fourth, we conduct a comprehensive evaluation on both transductive (seen diseases) and inductive (unseen diseases) splits, achieving a 5.4-fold improvement in ranking quality over baselines for novel diseases, highlighting the power of ontology-informed generalization. To our knowledge, this represents the first AI-driven ranker for disease-adjuvant recommendation, providing a concrete example of how AI can unlock the predictive potential latent within community-built ontologies.

Based on a comprehensive search of PubMed, bioRxiv, and Google Scholar (2020–2025), this work is, to our knowledge, the first to directly address the challenge of *de novo* adjuvant ranking for a given disease using ontology-grounded graphs. While prior work has focused on complementary problems such as antigen discovery (e.g., Vaxign/Vaxign2) or peptide adjuvant design (e.g., VaxinPAD), our approach is novel in its disease-first orientation. We introduce an AI-driven ranker that, given a disease, recommends a ranked list of suitable adjuvants. This represents a significant departure from traditional methods that often evaluate adjuvants in isolation.

### 4.1. Limitations

Several limitations exist. First, this study used incomplete data. Known disease–adjuvant links are sparse and uneven across disease families; negatives are often “unknowns,” not true non-associations. Knowledge bases are evolving and may lag behind the literature. Although VO, VIOLIN, and VaxKG exist to reduce such gaps, coverage remains imperfect. Second, we face challenges in ontology alignment and drift. Mapping adjuvant names across VO, VAC, and published papers is prone to error; updates to VO/VAC can alter identifiers or hierarchies, thereby affecting reproducibility. Third, our model currently lacks adverse event and administration awareness. The scorer does not yet penalize candidates using structured adverse-event semantics or route/dose constraints, which are crucial for translational use. The Ontology of Adverse Events (OAE) provides a formal scaffold we have not yet fully leveraged. Finally, evaluation limitations remain: macro-averaging over diseases can hide skew; bootstrap confidence intervals widen with small *n*; and text-similarity baselines understate the headroom relative to stronger retrieval-augmented systems.

### 4.2. Future Directions

We identify several key avenues for further work.

**(D1) Data enrichment:** We will curate more data from VO-linked sources by integrating VaxKG/VIOLIN graph exports and Vaccine Adjuvant Compendium (VAC) entries—all of which reference VO identifiers—so that each adjuvant candidate carries standardized attributes such as class, formulation, platform usage, and development stage. Concretely, we will build a VO-anchored schema (disease, vaccine, adjuvant, platform) and import releases with versioned IRIs for reproducibility. Nodes will be enriched with curated properties (e.g., Th1/Th2 skew, TLR targets, and known AE flags) to improve supervision and type-consistent negative sampling. Small, versioned graph slices (in JSON-LD/RDF format) will be published to make training and evaluation sets reusable.

**Furthermore**, we will test novel methodology for **knowledge-graph–aware ranking**. We aim to move from feature engineering to end-to-end learning on a VO-anchored graph that links diseases, pathogens, vaccine platforms, adjuvants, and, optionally, adverse events. Methodologically, the graph will be encoded using relational GNNs (e.g., R-GCN or HGT) or knowledge-graph embeddings (e.g., RotatE or ComplEx) and trained with a pairwise ranking loss for disease→adjuvant prediction. We will fuse unstructured evidence through contrastive retrieval (titles, abstracts, and labels), allowing text to serve as a first-class learning signal rather than a post-hoc feature. Lightweight ontology constraints—including relation typing, domain/range restrictions, and transitive closures—will be imposed during training and inference to reduce invalid edges.

For **uncertainty and calibration** in decision support, we will augment the ranker with calibrated uncertainty estimates so outputs are actionable in downstream triage. Specifically, we will report reliability diagrams and Expected Calibration Error (ECE) for top-*k* scores, and when appropriate, optimize with calibration-aware objectives. We also plan to deploy conformal prediction to return sets of adjuvants at target coverage 1 − *α*, enabling controllable recall with valid finite-sample guarantees. Finally, we will explore risk-aware ranking that penalizes candidates with high predicted adverse-event risk or administration mismatches, allowing explicit trade-offs between efficacy and safety.

## 5. Acknowledgments

We thank the developers of the VaxKG project for their foundational work and for insightful discussions. We are also grateful to the creators and maintainers of the Vaccine Ontology (VO), the VIOLIN knowledgebase, and the Vaxjo database for making their resources publicly available. YH was supported by NIH grant (U24AI171008).

## Declaration on Generative AI

The authors have not employed any Generative AI tools.

## Notes

### Competing Interest Statement

The authors have declared no competing interest.

### Summary of Updates

change of tilte to make it consistent with the initial paper title

